# Robust Detection of Brain Stimulation Artifacts in iEEG Using Autoencoder-Generated Signals and ResNet Classification

**DOI:** 10.1101/2024.09.30.615930

**Authors:** Jeremy Saal, Ankit N. Khambhati, Edward F. Chang, Prasad Shirvalkar

## Abstract

**Background:** Intracranial EEG (iEEG) is crucial for understanding brain function, but stimulation-induced noise complicates data interpretation. Traditional artifact detection methods require manual user input or struggle with noise variability, especially with limited labeled data.

**Objective:** We developed a supervised method to automatically detect stimulation-induced noise in human iEEG recordings using synthetic data generated by Variational Autoencoders (VAEs) to train a ResNet-18 classifier.

**Methods:** Multi-lead iEEG data were collected, preprocessed, and used to train VAEs for generating synthetic clean and noisy signals. The ResNet-18 model was trained on images of spectra generated from these synthetic signals and validated on real iEEG data from five participants.

**Results:** The classifier, trained exclusively on synthetic data, demonstrated high accuracy, precision, and recall when applied to real iEEG recordings, with AUC values greater than 0.99 across all participants.

**Conclusion:** We present a novel approach to effectively detect stimulation-induced noise in iEEG, offering a robust solution for improving data interpretation in scenarios with limited labeled data. Additionally, the pre-trained ResNet-18 model is available for the community to use, facilitating further research and application in similar datasets.

## Introduction

Intracranial electroencephalography (iEEG) recording techniques such as stereo-electroencephalography (sEEG) and electrocorticography (ECoG) provide invaluable insights into brain function ^1^. Recent studies have increasingly utilized iEEG electrodes for both recording neural activity and delivering electrical brain stimulation^2–4^. The advent of subacute, continuous iEEG recordings has opened new avenues for long-term monitoring of brain activity, enabling researchers to address important scientific questions^5–7^. Despite the advantages of intracranial recordings, they are still susceptible to various artifacts such as noise generated by brain stimulation (Fig. 1). This stimulation noise poses a significant challenge for accurate data interpretation, necessitating the development of robust detection methods to either exclude or further process noisy segments.

**Figure 1:**
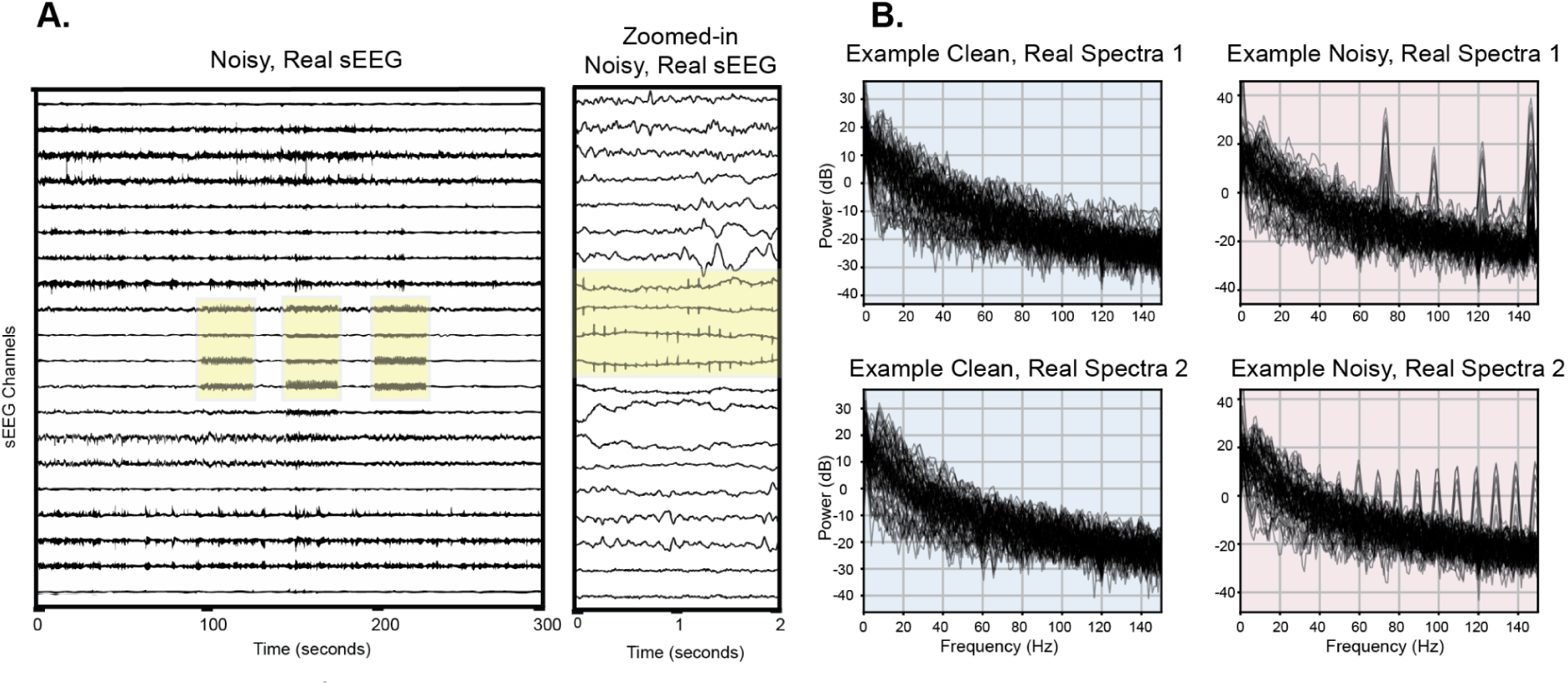
Example stereo-EEG with stimulation artifacts. (A) Time domain example of stimulation artifacts with artifacts highlighted in yellow. (B) Example clean, artifact-free data and noisy data with stimulation artifacts. Spectra from all channels are overlaid on top of each other.

Traditional methods for artifact detection, such as manual inspection and basic amplitude-based rejection based on the raw signal, are labor-intensive and often inadequate for addressing the complexity and variability of stimulation-induced noise in long-term intracranial EEG (iEEG) recordings. Even when stimulation is applied to a single bipolar pair of contacts among hundreds distributed across the brain, the resulting noise can spread across multiple channels. This widespread noise complicates the process of accurately isolating and removing artifacts, as the noise characteristics can vary significantly depending on electrode location and stimulation parameters. Consequently, basic detection methods may either fail to capture all instances of noise or mistakenly classify genuine neural activity as an artifact, compromising data integrity. Recent advances in deep learning, particularly the use of Convolutional Neural Networks (CNNs), have shown promise in automatically detecting artifacts in neural signals like EEG, MEG, and local field potentials (LFPs) ^8–11^. CNNs can improve noise detection accuracy as they excel at extracting complex patterns from time series data indicative of artifacts. However, such methods often require large, well-labeled datasets for effective training which are difficult to obtain in clinical settings. Although transfer learning has been explored as a solution to reduce the dependency on large datasets, it still typically requires some retraining, posing challenges when data is limited or when stimulation conditions are highly variable^8^. This highlights the need for an iEEG-based artifact detection system that can minimize or eliminate the need for extensive retraining, enhancing its applicability across diverse clinical and research scenarios.

Recent advancements in data augmentation techniques, which involve artificially increasing the diversity of a dataset through transformations such as rotation, scaling, or generating synthetic data, have improved the performance of deep learning models, particularly in situations with limited data availability^12^. However, there remains a significant gap in applying these methods specifically to stimulation-induced noise in iEEG recordings. Variational Autoencoders (VAEs), a type of generative data augmentation model, offer a powerful approach to address this gap. Generative models like VAEs are designed to learn the underlying distribution of input data to facilitate the generation of new, synthetic data that closely resembles the original signal. This synthetic data can effectively augment limited datasets, making it particularly valuable for training noise detection classifiers when labeled data is scarce. Some studies have begun to explore the use of other generative models, such as the transformer-based GAN introduced by Wu et al. (2024), to enhance classifier performance^13^; however, these approaches have yet to be fully integrated into noise detection systems. Moreover, the challenge of generating synthetic data that not only augments limited datasets but also accurately mimics the specific types of stimulation-induced artifacts found in iEEG recordings has not been fully addressed. To our knowledge, no prior studies have tackled the need for effective stimulation artifact detection in iEEG signals. This underscores the necessity for innovative approaches that leverage generative models like VAEs to create realistic, artifact-rich synthetic datasets, thereby improving the robustness and generalizability of noise detection models in iEEG recordings.

To address these challenges, we developed a novel approach for detecting stimulation-induced noise epochs in iEEG recordings by leveraging VAEs to generate synthetic iEEG data (Fig. 2). We then trained a ResNet-18 classifier—a deep convolutional neural network architecture known for its ability to extract complex patterns from images—on this synthetic dataset. Although ResNet-18 is typically used for image classification tasks, its residual learning framework adapts well to our noise detection task in neural data due to its capacity for capturing intricate patterns. Specifically, to leverage the benefits of CNNs and the ResNet-18 model, we used images of synthetic spectra as the input to the model rather than directly using neural signals. We validated our approach on real iEEG data from five individuals, demonstrating that the ResNet-18 classifier trained on synthetic data can distinguish between clean and noisy signal epochs. The classifier achieved outstanding accuracy, precision, and recall across diverse recording conditions, highlighting its potential as a reliable tool for improving the interpretability of iEEG data in both clinical and research settings. Our method effectively tackles the issue of limited labeled datasets and enhances the robustness of noise detection in long-term iEEG recordings. We have made available the fully trained model and code for easy use by the community (https://github.com/jersaal/ieeg-noise-detection).

**Figure 2:**
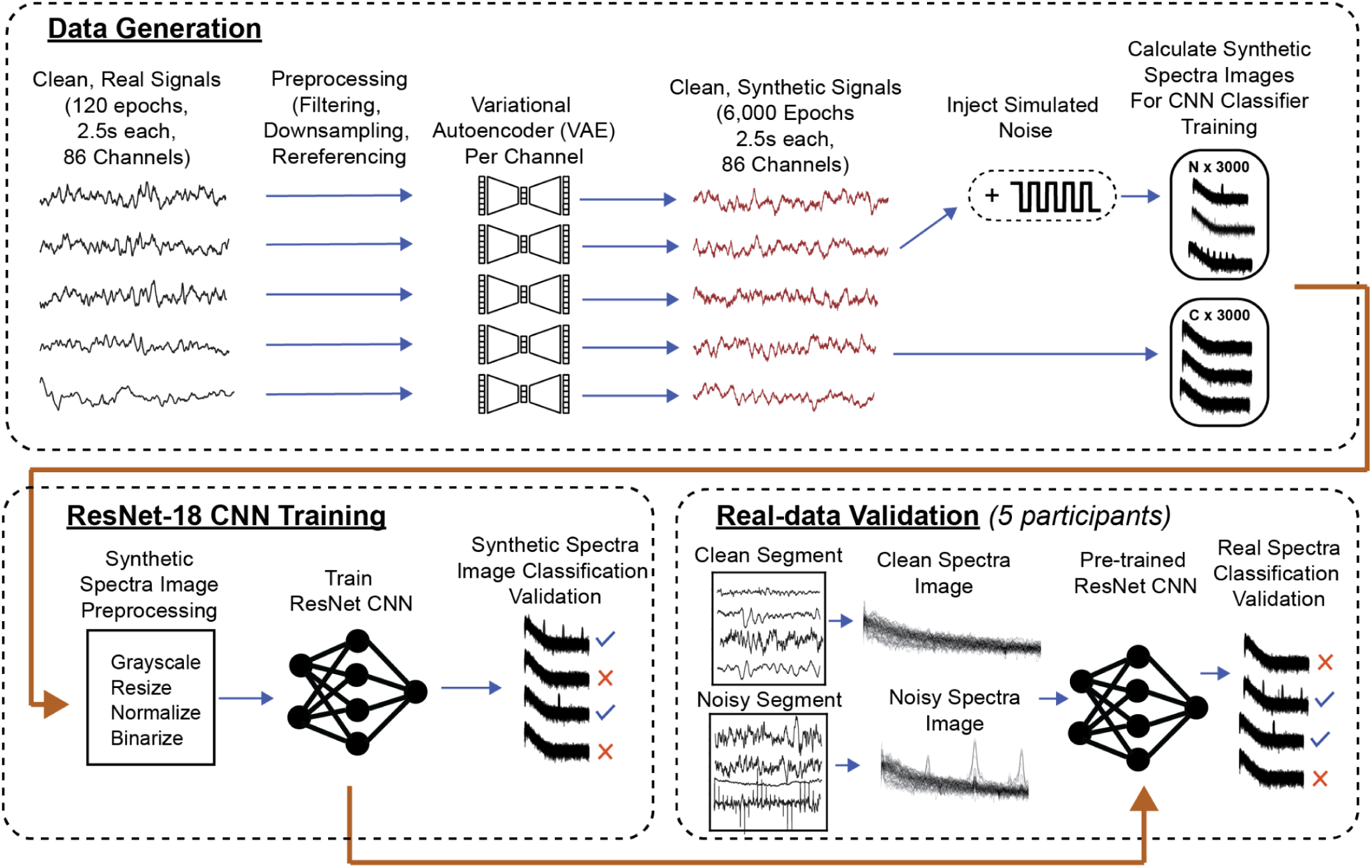
Data processing, model training, and validation pipeline. (Top) Data generation begins with clean, real iEEG signals (120 epochs, 2.5s each, 86 channels), which undergo preprocessing (filtering, downsampling, re-referencing) before being passed through a Variational Autoencoder (VAE) to generate synthetic clean signals. Simulated noise is injected into these synthetic signals to create noisy epochs, resulting in 3,000 clean and 3,000 noisy spectra images. (Bottom left) ResNet-18 CNN training pipeline involves preprocessing the spectra images (grayscale conversion, resizing, normalization, binarization), followed by training the ResNet-18 model to classify clean and noisy signals using synthetic data. (Bottom right) Real-data validation using five participants’ iEEG data involves applying the pre-trained ResNet-18 model to classify clean and noisy segments from real iEEG recordings, validated with spectra images generated from these recordings.

## Methods

We developed a comprehensive pipeline designed to detect stimulation-induced noise in iEEG signals. Our approach begins with the collection and preprocessing of iEEG data, followed by the use of VAEs to generate synthetic datasets that mimic both clean and noisy signals. These synthetic datasets are then used to generate images of multi-channel spectra which are then used to train a ResNet-18 classifier. To evaluate the robustness and generalizability of our approach, the trained classifier is validated using real iEEG data from five participants.

### Data Collection and Preprocessing

Data were recorded from five participants, each implanted with eight sEEG depth electrodes (PMT model 2102-16-091, Adtech models SD08R-SP05X-000 SD12R-SP05X-000 SD14R-SP05X-000) and up to two ECoG electrode arrays (PMT model 2110TX-16-076) over the sensorimotor cortex. The sEEG depth electrodes, containing 8-14 contacts spaced 5 mm apart, targeted regions including the bilateral anterior cingulate cortex (ACC), insula, subgenual cingulate (SGC), ventral thalamus, ventral striatum, and periventricular gray (PVG). Electrode placement was guided by BrainLab iPlan and Elements software, and recordings were captured using a 256-channel Nihon Kohden system (NK model PE-201A). Data were sampled at either 2,000 Hz or 10,000 Hz. Laplacian re-referencing was applied to enhance spatial resolution and reduce common noise across neighboring electrodes; this referencing scheme was also chosen to match the referencing scheme of future analyses. Subsequently, data were notch filtered at 60Hz and the harmonics of 60Hz, high-pass filtered from .1 Hz, and low-pass filtered under 250 Hz. After re-referencing and filtering, data were downsampled to 1,000 Hz. Recorded signals contained numerous periods of bipolar stimulation at adjacent sEEG contacts using biphasic pulses with the following range of parameters: frequency 5-150Hz, pulse width 100-400 us, amplitude 0.5-6mA, duration 5-300s.

### Variational Autoencoder Architecture

We used a Variational Autoencoder to generate synthetic iEEG data tailored specifically for noise classification using the Python programming language and the PyTorch machine learning library. We trained separate VAE models for each iEEG channel and ensured that the generated synthetic data closely matched the unique characteristics of each channel by comparing the autocorrelation functions.

The VAE architecture consisted of an encoder and a decoder, both implemented using 1D convolutional layers to capture the temporal structure of the iEEG signals. The encoder comprised three 1D convolutional layers with rectified linear unit (ReLU) activations, progressively increasing the number of filters (128, 256, 512) to extract increasingly complex features from the input signals. The kernel size (15), stride (1), and padding (3) were selected based on the intrinsic timescale of expected changes in the stereo EEG signal, corresponding to 15 milliseconds. After the convolutional layers, the output was flattened and passed through two fully connected layers to compute the mean and log-variance of the latent space distribution. We used 50 latent factors to capture the essential features of the input data, balancing the model’s capacity to represent complex temporal patterns in the iEEG signals with the need to avoid overfitting. This number was selected based on preliminary experiments that indicated it provided sufficient dimensionality to capture the key variability in the data without introducing excessive complexity that could lead to overfitting or increased computational burden. The VAE decoder mirrored the encoder’s architecture, using transposed convolutional layers to reconstruct the LFP signals from the latent space representation.

For training the VAE, we used data from one participant (P02) covering 86 channels. Ten 30-second segments (5 minutes total) of visually-identified, stimulation-artifact-free data were randomly selected from recordings over the course of ten days to capture different periods of neural activity. This selection process ensured that the data represented various times of day. We chose five minutes of data to demonstrate that our method can effectively work with minimal input. The 30-second segments were further divided into 2.5-second windows. Each window was normalized by Z-scoring the raw iEEG voltage values across channels to ensure comparable amplitudes. The VAE-generated synthetic data were subsequently used to train a ResNet-18 classifier.

### Hyperparameter Tuning and Training

VAE training was optimized by minimizing a combination of reconstruction loss and Kullback-Leibler (KL) divergence. The reconstruction loss was calculated using mean squared error (MSE) between the original and reconstructed signals, while the KL divergence regularized the latent space, encouraging meaningful and smooth representations. The learning rate was set to 0.0001, and the model was trained with a batch size of 5 time-windows per batch. The model was trained separately for each channel, with the data split into training (80%) and validation (20%) sets. Early stopping was employed to prevent overfitting, based on validation loss, with the training process monitored over 50 epochs. The final model was selected based on the lowest validation MSE.

### Synthetic Data Generation

Once trained, VAEs were used to generate synthetic iEEG data separately for each of the 86 channels. For each individual channel, we generated 6,000 synthetic clean epochs spanning 2.5 seconds. Of these epochs, 3,000 will have simulated noise added. The synthetic data was generated by drawing random points from the learned latent space of the VAE, where the latent space represents a compressed version of the original iEEG data. These points were then passed through the decoder to reconstruct synthetic iEEG signals. This process allows for the creation of entirely new signal epochs that capture the characteristics of the original data without being direct replicas.

### Verification of Synthetic Data Using Autocorrelation Analysis

To assess whether the synthesized iEEG time series reflected the temporal characteristics of real iEEG signals, we evaluated autocorrelations of clean, synthetic data. Autocorrelations capture the periodic nature of the signals, which vary by frequency and exhibit distinct 1/f characteristics. For this analysis, we compared five minutes of real iEEG data to five minutes of synthetic data, each consisting of 120 epochs of 2.5-second windows. The autocorrelation function (ACF) was calculated for each epoch within each channel; ACFs were averaged across trials to produce a mean ACF for both real and synthetic data. We assessed the similarity between real and synthetic iEEG signals by calculating Pearson’s correlation coefficients between the mean ACF curves for each channel. Statistical significance of correlations across all channels was evaluated using Fisher’s combined probability test^14^.

### Simulated Noise Addition

To simulate stimulation artifact noise in the synthetic iEEG data, square wave signals of various amplitudes and frequencies were added to a subset of channels to replicate the stimulation patterns commonly used in electrical brain stimulation. Synthetic noise was introduced within the 3,000 synthetic clean iEEG epochs. The noise amplitudes were selected from the set [4, 6, 10, 14 arbitrary units], and the frequencies ranged from 5 Hz to 140 Hz in 5 Hz increments, with an additional low-frequency component at 0.5 Hz to cover the lower spectrum of potential noise. The noise amplitudes were chosen to be proportional to the background signal amplitude, ensuring that the injected noise realistically reflects scenarios where stimulation artifacts have a comparable or higher amplitude than the underlying neural signals. To further simulate realistic noise patterns, the amplitude of the noise was scaled inversely with frequency using a linear relationship, meaning that lower frequencies had higher amplitudes. For example, a 5 Hz noise component would have a greater amplitude than a 140 Hz component, with noise amplitude decreasing as frequency increased. The number of noisy channels per epoch varied between 3 and 11; we also added noise signals at multiple harmonics (ranging from 1 to 15) of the base frequency to resemble complex noise patterns that mimic real data.

### Power Spectral Density Calculation and Image Generation

The power spectral density (PSD) of the iEEG signals was calculated for each epoch using Welch’s method, with a segment length of 1,024 samples. PSDs were chosen because they provide a detailed representation of the frequency content of the signals, which is crucial for distinguishing between clean and noisy epochs. PSDs, representing both the original and synthetic signals, were plotted on a logarithmic (base 10) scale, with the y-axis range set to the minimum and maximum log-transformed power values observed in the data. Frequencies ranged from 1 to 150 Hz. The resulting plots were saved as PNG images with a resolution of 100 dpi and were subsequently used as inputs to the ResNet classifier to classify clean and noisy signals.

### ResNet Model Training and Configuration for Noise Classification

To classify whether a signal was noisy or clean, we used a ResNet-18 classifier, chosen for its robust architecture and its ability to capture complex patterns in image data through its deep residual learning framework^15^. ResNet, short for Residual Network, is a type of convolutional neural network (CNN) designed to address the vanishing gradient problem that can occur in deep networks. It achieves this by introducing “skip connections,” which allow the network to bypass certain layers, making it easier to train very deep networks by ensuring gradients can flow directly through these connections. The ResNet-18 model specifically consists of 18 layers and is known for its efficiency in feature extraction and high performance in various classification tasks. The ResNet-18 model, initially pre-trained on the ImageNet dataset, was trained on 3,000 clean and 3,000 noisy spectra images and class labels, calculated from synthetic iEEG signals per participant.

To optimize the model’s performance, we applied several preprocessing steps to simplify the images. First, the images were converted to grayscale. Next, images were resized to 672×672 pixels to match the input dimensions required by the model. Each image’s pixel values were then normalized to have a mean of 0.5 and a standard deviation of 0.5, ensuring consistent pixel ranges for stable and efficient training. Finally, the images were binarized using a consistent threshold of 0.8 for all images, determined by visual inspection. During this process, we ensured that all signal peaks were preserved, enhancing contrast by setting pixel values above the threshold to black and those below to white.

We modified the model’s initial convolutional layer to accept single-channel grayscale images (by changing the input channels from 3 to 1) and replaced the final fully connected layer to output two logits (for binary classification). These logits were then converted to probabilities using the softmax function^16^. The output probability represents the likelihood that an input image belongs to the ‘noisy’ class (class 1), with the probability of the ‘clean’ class (class 0) being its complement. Using five-fold cross-validation, the dataset was split into different training and validation subsets for each fold. Models were optimized using cross-entropy loss, a common objective function for binary classification, and the Adam optimizer, which adapts the learning rate during training. The learning rate was set to 0.0001, and models were trained across a maximum 20 epochs for each fold, with early stopping if validation loss did not increase for 5 consecutive epochs. To evaluate the performance of the model in distinguishing between synthetic clean and noisy signals, we computed the area under the Receiver Operating Characteristic curve (AUC-ROC), accuracy, precision, and recall.

### Validation Using Real Patient Data

To evaluate the generalizability and robustness of the ResNet-18 model, we conducted a validation analysis using real iEEG data from five patients undergoing sEEG based brain stimulation and recording. For validation of the performance on real-world data, the ResNet-18 model was trained on all of the synthetic data (i.e., instead of using 5-fold cross-validation). We set a decision threshold of 0.2 to ensure that noisy signals are more sensitively identified. This threshold was conservatively chosen to address the challenge of misclassifying real signals as noise, which is critical for maintaining the integrity of the clean segments. While the VAE was trained on real iEEG data from one participant, the ResNet classifier was trained exclusively on synthetic iEEG data, with no exposure to real patient data during its training phase. This validation rigorously assessed the model’s ability to generalize from synthetic training data to real-world iEEG data.

For each participant, real iEEG data underwent preprocessing consistent with the initial preparation steps, including segmentation into 2.5-second windows surrounding the stimulation onset. These windows were categorized as either “clean” (from -5 to -2.5 seconds prior to stimulation onset) or “noisy” (from 0 to 2.5 seconds after stimulation onset) based on stimulation timestamps acquired from our stimulation system. Image generation and pre-processing were identical to that used for the synthetic data. The trained ResNet-18 model was then applied directly to these spectra images to classify each epoch as either noisy (1) or clean (0). The model’s performance was evaluated using accuracy and the area under the curve (AUC) of the receiver operating characteristic (ROC), precision, recall, and F1 score. We conducted permutation testing to assess the statistical significance of the observed AUC values. Permutation testing involved randomly shuffling the labels of the validation data 1,000 times to generate a distribution of AUCs under the null hypothesis, where no true association exists between the spectra and their labels. The p-value was then calculated by comparing the actual AUC to this distribution, providing a robust measure of the model’s classification performance relative to random chance. This permutation-based approach offers a stringent test of the model’s ability to generalize to real-world conditions, ensuring that the high AUC values observed are not simply due to overfitting or random correlations in the data.

## Results

### VAE Training and Validation

We used per-channel Variational Autoencoders (VAEs) to generate 6,000 clean iEEG epochs tailored for classification, as our dataset contained limited labeled data, 3,000 of which subsequently had simulated noise added. The VAE models were initially trained on a dataset comprising 10 stim-artifact-free iEEG recordings, each 30 seconds in duration (5 minutes total) from participant P02. To avoid overfitting, we employed early stopping based on validation loss (Fig. 3A, bottom right). Example traces for real and synthetic iEEG can be seen in Figure 3A (top left and top right). After adding synthetic noise to 3,000 of the generated epochs, the noise simulation method produced clear PSD peaks at the added noise frequencies (Fig. 3C).

**Figure 3:**
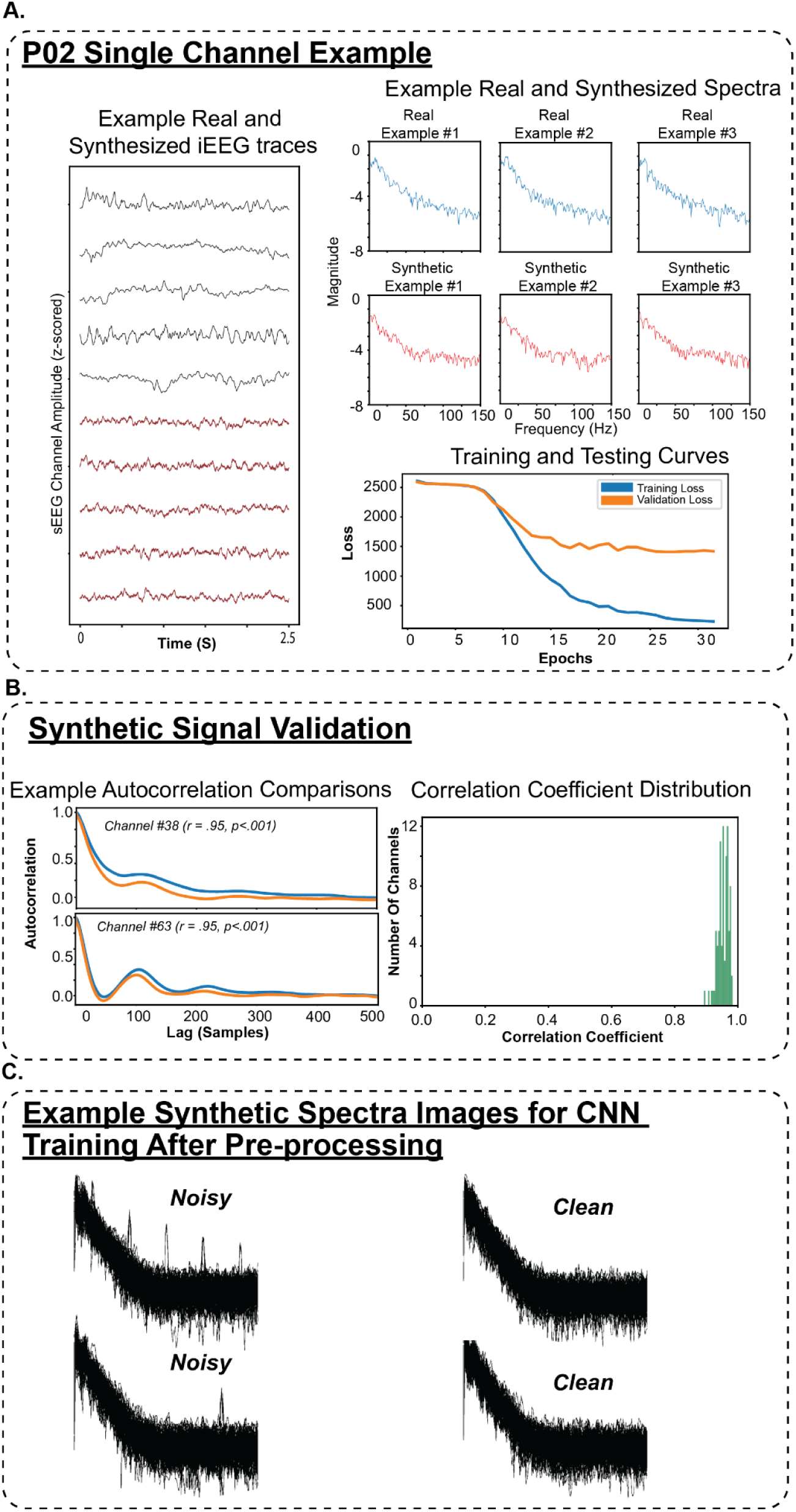
Generation and Validation of Synthetic iEEG Data Using a Variational Autoencoder (VAE). (A) Example of real and synthetic sEEG traces from Channel 7 of participant P02, showing that the VAE accurately replicates the temporal dynamics of real signals. Spectral comparisons between real and synthetic signals (right) demonstrate that the synthetic signals preserve the frequency content of the real data across various examples. The training and testing loss curves indicate stable training performance after 37 epochs. (B) Validation of the synthetic signals using autocorrelation comparisons between real and synthetic data for two channels, showing a strong match between their temporal structure across the first 500 samples. (C) Example of pre-processed spectra images for CNN training, illustrating noisy and clean synthetic iEEG signals.

### Synthetic Data Verification

After training, we assessed the fidelity of the synthetic iEEG data by comparing it to the original real iEEG signals using Pearson’s correlation between their respective autocorrelation function (ACF) curves. Pearson correlation coefficients were exceptionally high, ranging from 0.91 to 0.99 across all channels (Fisher’s combined probability p<.001), confirming that the synthetic data accurately captured the temporal structure of the real iEEG signals (Fig. 3B).

### ResNet Model Performance With Synthetic iEEG Data

The ResNet-18 model, used for binary classification of noisy versus clean iEEG signals, was trained exclusively on synthetic data generated by the VAE models. Using five-fold cross-validation, the ResNet-18 model achieved F1, precision, and recall of 99.95%, 99.93%, and 99.96%, respectively. AUC-ROC was greater than 0.99, indicating outstanding performance in discriminating synthetic clean and noisy signals (Fig. 4A)

**Figure 4:**
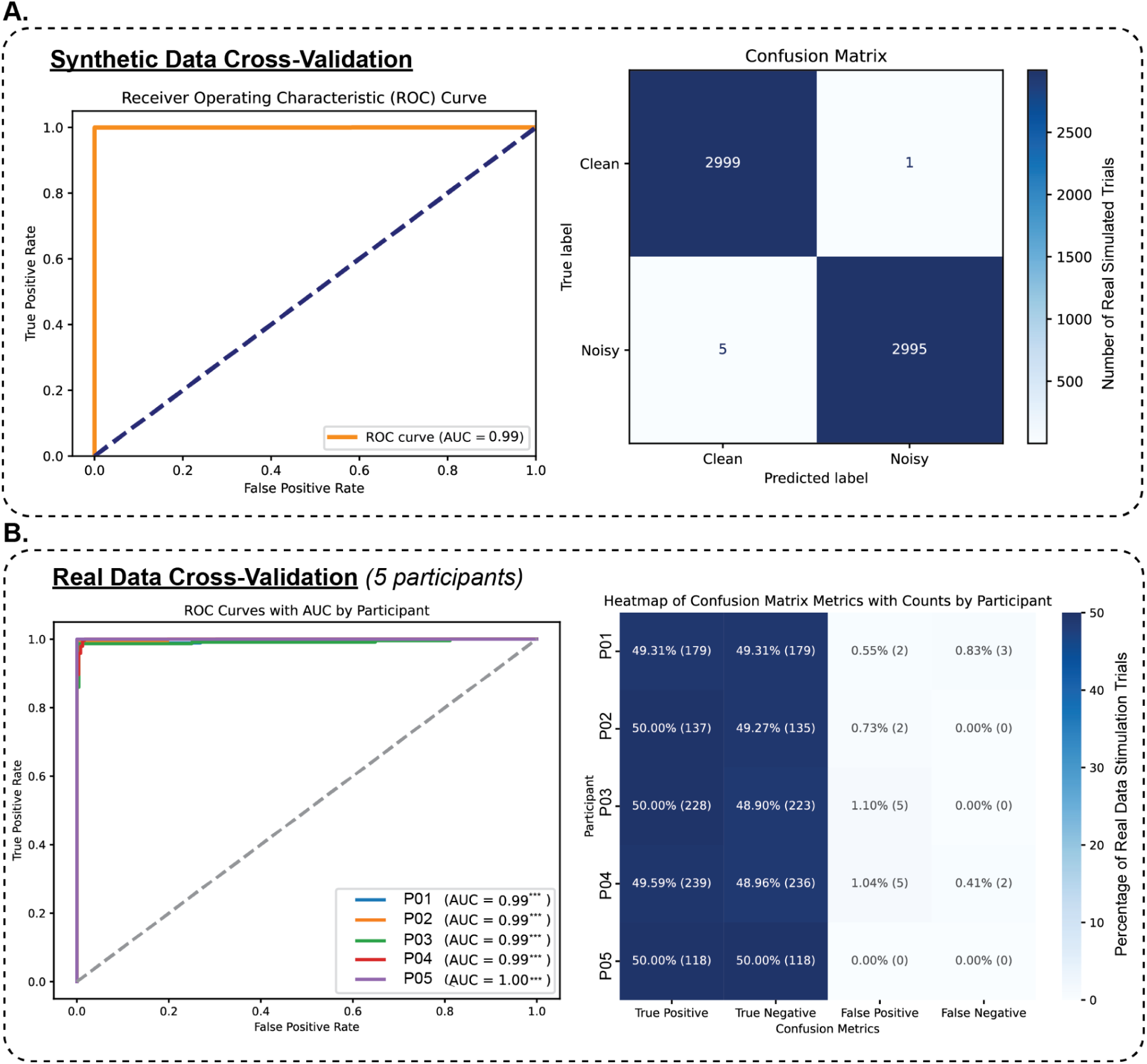
Classifier performance on synthetic and real data. (A) Synthetic Data Cross-Validation: Features an ROC curve with an AUC of 1.00, indicating near-perfect classifier performance on synthetic data, and a confusion matrix showing no misclassifications, highlighting the model’s accuracy in idealized conditions. (B) Real Data Cross-Validation (5 participants): ROC curves for each of the five participants with AUC scores close to 1.00, demonstrating the classifier’s near-perfect performance across diverse real patient data, and a heatmap of confusion matrices detailing performance metrics for each participant. P02 was used for training the VAE.

### Stimulation Noise Classification Validation Across Participants

To validate real-world generalizability of the trained ResNet-18 noise classifier model, we performed validation on five real iEEG datasets derived from different participants (P01, P02, P03, P04, P05) each with 363, 274, 456, 482, and 236 stimulation epochs, respectively. These datasets were not used during training and represent both diverse stimulation parameters and recording conditions. The ResNet-18 model again demonstrated outstanding performance, achieving classification accuracies of 98.35%, 99.64%, 98.68%, 98.34%, and 100.00% for participants P01 through P05, respectively (Fig. 4B). AUC-ROC values were equally good measuring 0.9968, 0.9986, 0.9919, 0.9993, and 1.0000 for P01-P05. Permutation testing, conducted with 1000 permutations for each participant, yielded statistically significant results across all datasets (p<.001). F1, Recall, and Precision scores are reported in Table 1. These results highlight the model’s robust sensitivity, reliability and generalizability in detecting stimulation noise in real-world iEEG signals across multiple participants. Furthermore, classification time per spectra image was 8 milliseconds on average, highlighting the potential for this approach to be used for real-time noise detection.

**Table 1:**
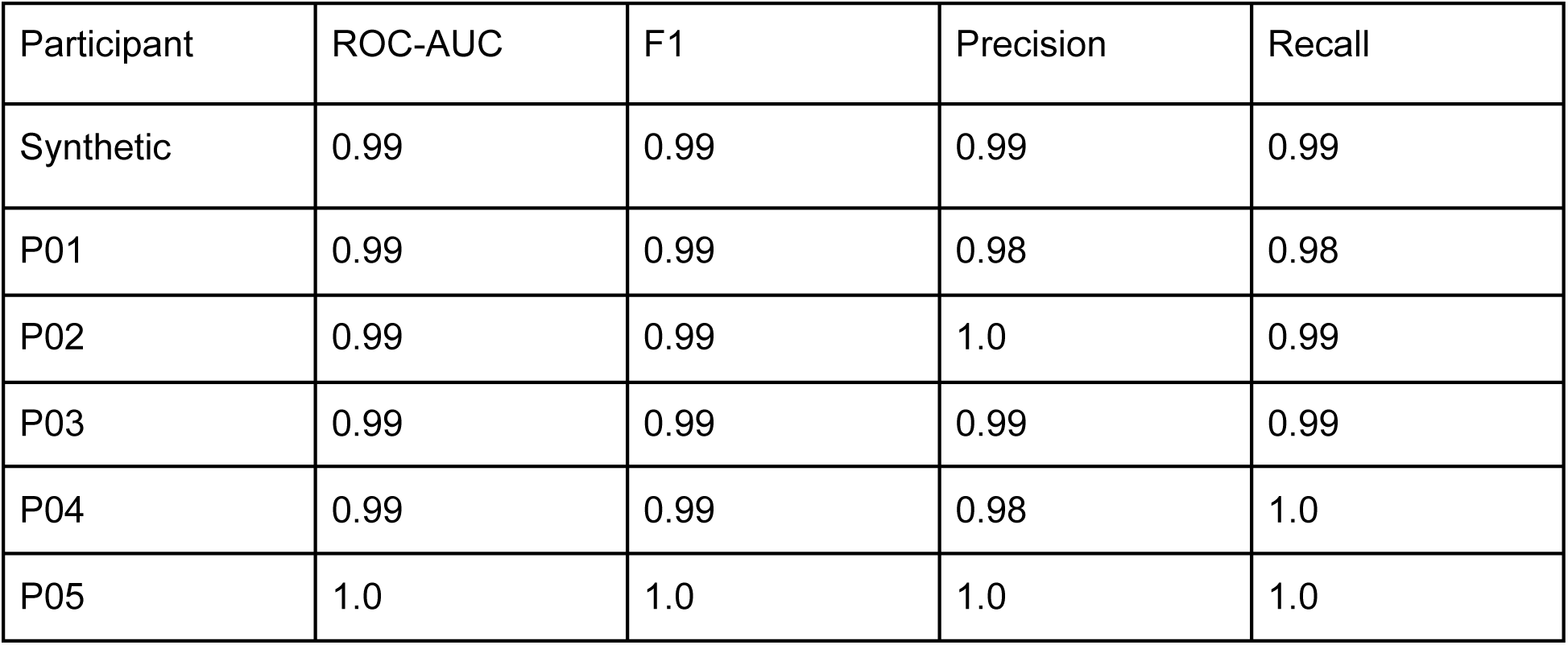
Performance Metrics for ResNet-18 Classifier in Detecting Stimulation-Induced Noise Across Participants. The table summarizes the classification performance of the ResNet-18 model for each participant (P01–P05) based on ROC-AUC, F1 score, Precision, and Recall. ROC-AUC values close to 1.00 indicate excellent model discrimination between noisy and clean signals. F1, Precision, and Recall scores are consistently high across participants.

| Participant | ROC-AUC | F1 | Precision | Recall |
| --- | --- | --- | --- | --- |
| Synthetic | 0.99 | 0.99 | 0.99 | 0.99 |
| P01 | 0.99 | 0.99 | 0.98 | 0.98 |
| P02 | 0.99 | 0.99 | 1.0 | 0.99 |
| P03 | 0.99 | 0.99 | 0.99 | 0.99 |
| P04 | 0.99 | 0.99 | 0.98 | 1.0 |
| P05 | 1.0 | 1.0 | 1.0 | 1.0 |

## Discussion

In this study, we developed and validated a novel approach for detecting stimulation-induced noise in iEEG recordings by leveraging synthetic data generated with Variational Autoencoders. Remarkably, the ResNet-18 classifier, trained exclusively on synthetic iEEG data, demonstrated strong generalizability to real iEEG data across multiple participants, achieving outstanding accuracy, precision, and recall. Our approach effectively addresses the challenges of automated noise detection in iEEG recordings, which holds significant potential, particularly in scenarios with limited labeled datasets. This performance further highlights the potential of VAEs for creating realistic synthetic neural signals that can serve as a robust foundation for training deep learning models in neural signal processing.

The ability of our model to accurately classify noise in iEEG data is crucial for enhancing the interpretability of recordings, especially in studies involving deep brain stimulation (DBS), where stimulation artifacts can confound analysis. Notably, the performance of our model, with F1 scores ranging from 0.97 to 1 across participants, compares favorably to other CNN-based approaches for artifact detection. For example, a pre-trained CNN model for general iEEG artifact detection, not just stimulation-induced artifacts, reported in a related study, achieved an F1 score of 0.89 when tested on held-out data^8^. However, it was only after fine-tuning the model with a small subset of the target dataset that the F1 score improved to 0.97, a result comparable to the performance achieved by our model. This highlights the need for dataset-specific adaptation in some models. In contrast, our model, trained solely on synthetic data, demonstrated high performance without the need for additional fine-tuning on real data, showcasing its strong generalizability across participants and recording conditions.

This has significant implications for noise detection in naturalistic datasets, as it demonstrates the potential to generate and train models on synthetic noise that closely mimics real-world artifacts, thereby reducing the need for extensive manually labeled datasets. By using raster images of PSDs rather than raw vectors or PSD values directly, our approach leverages the spatial patterns in these images, which may capture complex noise characteristics more effectively than time series signals. Furthermore, this approach does not require a priori knowledge of the stimulation frequency used. This novel use of visual representations for classification suggests that noise sources in iEEG data can be effectively identified and characterized based on their visual patterns, offering a promising new direction for artifact detection across diverse recording environments.

Despite these promising results, there are limitations to our study that warrant further exploration. It is important to note that our method focuses on detecting and excluding noisy epochs rather than removing artifacts, as our goal is to ensure that non-stimulation periods are as clean as possible for accurate data analysis. Our current study was conducted on patients without epilepsy, and while this provided valuable insights, future work should extend this approach to patients with epilepsy to assess its generalizability in clinical settings where seizure activity and its associated artifacts may be more prevalent. Moreover, this approach could be extended to detect artifacts arising from variable sources such as movement or electrical interference. A key strength of our study is the inclusion of both cortical and subcortical targets, providing generalization across the brain. By necessity, electrode trajectories resulted in the oversampling of some regions, which could introduce bias when analyzing unusual electrode montages. Further, the pattern of noise injected into the synthetic data may not fully replicate real-world conditions where volume conduction from stimulation might create more noise in nearby contacts than in distant ones. Additionally, our model’s fast classification speed indicates that it could be applied for real-time noise detection in clinical environments, future research should be conducted to assess real-time implementation including image generation.

Overall, our findings highlight the potential for integrating advanced noise simulation with synthetic data generation to improve the detection and interpretation of artifacts in iEEG recordings across diverse clinical and research contexts. Code as well as a pre-trained ResNet classifier are available at https://github.com/jersaal/ieeg-noise-detection.

